# Pathogen Population Structure Can Explain Hospital Outbreaks

**DOI:** 10.1101/210534

**Authors:** Fabrizio Spagnolo, Pierre Cristofari, Nicholas P. Tatonetti, Lev R. Ginzburg, Daniel E. Dykhuizen

## Abstract

**Objective:** To analyze Hospital Acquired Infection (HAI) outbreaks using microbial population biology dynamics in order to understand outbreaks as a biological system.

**Design:** Computational modeling study.

**Methods:** The majority of HAI transmission models describe dynamics on the level of the host rather than on the level of the pathogens themselves. Accordingly, epidemiologists often cannot complete transmission chains without direct evidence of either host-host contact or large reservoir populations. Here, we propose an ecology-based model to explain the transmission of pathogens in hospitals. The model is based upon metapopulation biology, which describes a group of interacting localized populations and island biogeography, which provides a basis for how pathogens may be moving between locales. Computational simulation trials are used to assess the applicability of the model.

**Results:** Results indicate that pathogens survive for extended periods without the need for large reservoirs by living in localized ephemeral populations while continuously transmitting pathogens to new seed populations. Computational simulations show small populations spending significant portions of time at sizes too small to be detected by most surveillance protocols. The number and type of these ephemeral populations enable the overall pathogen population to be sustained.

**Conclusions:** By modeling hospital pathogens as a metapopulation, observations characteristic of hospital acquired infection outbreaks for which there has previously been no sufficient biological explanation, including how and why empirically successful interventions work, can now be accounted for using population dynamic hypotheses. Epidemiological links between temporally isolated outbreaks are explained via pathogen population dynamics and potential outbreak intervention targets are identified.

## Introduction

Hospital acquired infections (HAIs) adversely impact patient care and outcomes. Worldwide, the rate of HAIs for patients in the intensive care unit approaches 30% and deaths due to antibiotic resistant pathogens are expected to outpace even cancer deaths and reach 10 million annually by 2050 ^1^. Central to the fight to lower the number of HAIs are mechanisms that lower or slow transmission of pathogens within hospital environments ^2^. Many of these mechanisms are neither novel nor highly technical. Patient isolation ^3^, patient ^4^ and environmental disinfection ^5^, as well as handwashing protocols ^6^, remain the most successful and cost effective means of controlling transmission of HAIs.

The application of genomic tools to hospital epidemiology has also become a valued outbreak control tool, enhancing both magnification and resolution into outbreaks ^7^. However, even as genomic analyses coupled with epidemiological data have provided a deeper understanding of how nosocomial infections spread in hospital outbreaks, there remains two interacting aspects of outbreaks where understanding remains lacking: the discontinuity in outbreak cases and the lack of known pathogen reservoirs. The discontinuity of infections in outbreaks are periods of time in which there is a gap between recognized cases. Often, additional infections are later identified as being caused by the same strain ^8–12^ without any epidemiological link between the infected hosts (such as asymptomatic carriers). Thus discontinuity can sometimes be partially explained by genomic data and surveillance monitoring during and after outbreaks, such as in the identification of previously unknown infections ^12^. Additionally, these discontinuous “breaks” can occur for unexpectedly long periods of time, during which hospital epidemiologists may believe that a particular pathogen is no longer present within their facility.

When new cases are identified after such breaks and the strain is matched to a previous outbreak strain, a reservoir population is presumed to exist ^2^. The location and size of the reservoir population is often never identified, however some assumptions about reservoirs are common. For instance, reservoirs are thought to be large populations ^13–15^, although there are cases in which populations of unknown but lesser size are called reservoirs ^2,16^. In each of these situations, however, the source population is considered to be one from which many potential infections can be founded. This presupposes that the reservoir is a source population, with individual infections being sink populations. Assumptions concerning such source-sinks may turn out to be incorrect.

Another assumption concerning reservoirs is that they are durable and exist for extended periods of time ^17,18^. Long term sources have been associated with infections such as Legionnaires Disease ^19,20^, which is often associated with bacterial contamination in water supply systems. Recent studies have begun to elucidate the role of the hospital environment on the existence and persistence of possible nosocomial reservoir populations ^21–24^. However, these environmental hospital populations violate the assumption that reservoirs are large. Further, the duration of these small populations, while longer than previously suspected, is by no means permanent ^25^.

This necessitates the possibility that hospital acquired infections can be caused by pathogens from these ephemeral environmental populations ^26^, even when these populations are small. Importantly, if a pathogen from one of these populations is capable of causing an infection in a susceptible host, then that pathogen is also capable of founding a new environmental population somewhere else within the hospital environment, with the size, growth rate, and duration of that population being dependent upon a series of independent factors, not the least of which is how well adapted the bacterial clone is to surviving under such conditions and how often that population’s location is cleaned by facility staff.

The resulting conditions are such that hospital pathogens can survive for prolonged periods of time by emigrating from one ephemeral population to another until one of three possible outcomes is reached: bacterial cells will eventually die off, be killed through the regular process of hospital cleaning, or cause a new infection. If the cells are able to infect a new host, the size of the infecting pathogen population will grow exponentially and many new populations can be founded, starting the process anew. The result is that the pathogen population survives as a *metapopulation* (Fig 1 A-B).

**Fig. 1.**
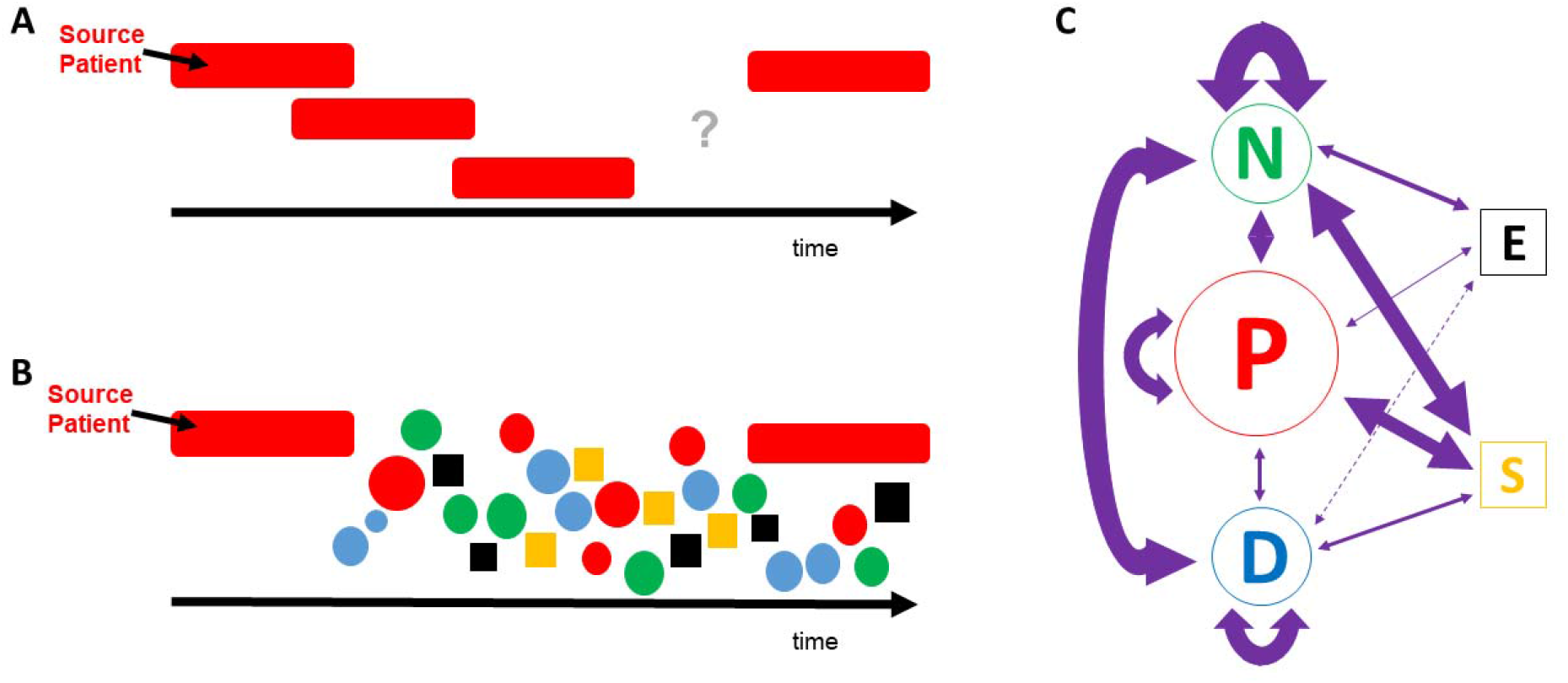
The Ephemeral Island Metapopulation Model in Hospital Environments. Colors represent each patch class: Patients in red; Nurses in green; Doctors in blue; Equipment in black; and Stationary surfaces are yellow. (A). A Gantt style diagram of a hospital outbreak with overlap of identified infected patients (elongated red bars) over time. The undefined area denoted by the question mark is a discontinuity in the epidemiological analysis of transmission typical of outbreaks. (B). A similar depiction of a hospital outbreak under the proposed model, where many ephemeral islands can bridge the discontinuity so long as the metapopulation exists within the hospital facility. (C). Graphic representation of transmission routes in the model. Transmission route arrows and patch areas represent (approximately) the strength of transmissions modeled. Transmission routes are in linear scale and patch classifications are in log scale (relative to each other). Dashed transmission line between Doctors and Equipment denotes low interaction.

Here, we propose a model for HAIs that is based upon the ecological concept of a metapopulation ^27^ and is further informed by island biogeography ^28^. Briefly, a metapopulation is defined as an assemblage of interacting populations ^29^. We are proposing that hospital pathogen metapopulations survive across a patchy distribution of habitable areas within a larger inhabitable landscape by existing in smaller populations for some period of time. Individuals from one “patch” or “island” move to other locales after some period of time before the resources required to survive in the previous patch are exhausted and the resident population goes extinct. Smaller populations exist within a particular hospitable patch for a period of time, but not indefinitely. The patches can vary spatially, like small islands in an inhospitable sea, or temporally, such as a blooming plant in a desert might for a group of insect nomads. Then, after some unknown amount of time has passed, individuals from an island population either die off (extinction) or move on to other islands, within which they can found a new island population or join one already existing. The amount of time that a population can stay on an island depends on factors that determine how long that island remains capable of supporting the population, as each island can only support a limited number of individuals. A biologically important characteristic of metapopulations is how well these smaller populations (or individuals from them) can move from island to island. In macroscopic species, this is a function of an organism’s ability to migrate, but for microbes the relevant mechanism is human-mediated carriage or transmission.

From some time, the applicability of metapopulations to problems of pest control ^30^, parasitism and infections ^31^ have been recognized. The further application of metapopulation ecology to nosocomial infection transmission should also be evident. However, instead of naturally imposed limits on patch viability, the likelihood of an island being sustainable or not is dictated by human behaviors such as hand washing, patient isolation, antibiotic stewardship protocols, healthcare worker cohorting, and the frequency of environmental cleaning ^32^. Not surprisingly, all of these interventions are thought to be effective infection control measures in hospital environments ^22,32–34^, albeit with an adequate science base capable of explaining why ^35,36^. The hypothesis proposed here provides a biological rationale for the empirical successes of these interventions by considering pathogens at the population level without reliance on classical assumptions. Understanding metapopulation dynamics of pathogens in this way has the potential to provide a wide scientific foundation upon which hospital epidemiology can be interpreted and further developed.

## Methods

Our model investigates the ability of pathogens to spread and survive via numerous interactions between different carriers. Using Monte Carlo simulation, we model the interactions between different islands and investigate the long term viability of the pathogen metapopulation. The model considers five classes of island/patch. Three of these are potential human hosts: patients (P), nurses (N), and physicians (D). Two account for the environment: mobile equipment (E) and non-mobile surfaces (S), modeling the role of the environment in the potential survival and transmission of HAI pathogens. Through these five classes most typical interactions can be simulated.

General properties of interactions for each island type can be listed as follows: (D) interact with all patients (P). (N) do not necessary interact with all (P), but the frequency of (N)-(P) interactions is greater than the frequency of (P)-(D). (S) do not directly interact with each other. (P) only have direct interaction with (E) and (S) in their immediate environment. (S) and (E) can be cleaned more effectively than (N), (P) and (D) (Fig 1C).

Several parameters are needed to simulate the pathogen population over the different islands. The maximum population size (carrying capacity) of a patch *Max_i_* corresponds to the maximum number of pathogens that the island of type *i* can support. The number of interactions, over a given period of time, between an island of type i and an island of type *j* is noted as *N_ij_*. The probability of pathogen transmission, i.e. the probability for the pathogen to be found on *j* after an interaction with i is *P_ij_*. The frequency of cleaning an island is *C_i_*. The efficacy of cleaning, i.e. the fraction of the patch population killed through cleaning is *L_i_.*Typical values for these parameters are given in Figure 2. The population, when left alone on an island, has a doubling time τ. Here, the value of τ is 8 hours.

**Fig. 2.**
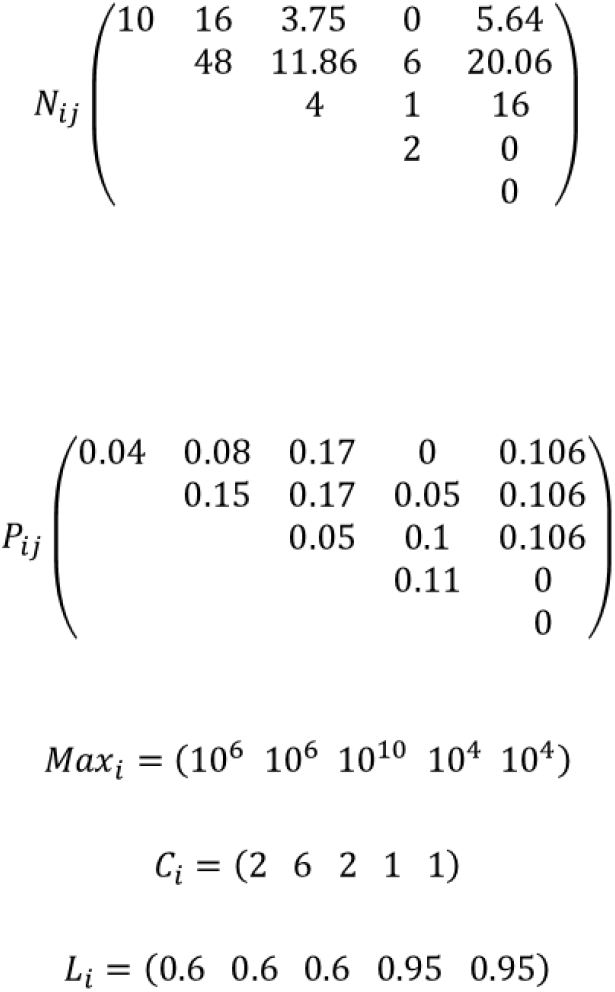
Parameter values used for the current simulations. The matrices *N_ij_* and *P_ij_* represent the number of interactions within and between parameter classes and the level of transfer of live bacterial cells during the interaction, respectively. The *N_ij_* matrix contains number of interactions per 8 hour shift and is based upon publicly available data sources for a range of health care facilities. The carrying capacity for any class of island is dependent upon patch type but is impacted by the frequency of cleaning (*C_i_*) as well as the thoroughness of that cleaning (*L_i_*). The regularity and thoroughness of cleaning is expected to keep the metapopulation from reaching equilibrium on a consistent basis, raising the overall importance of the level of detectability of a pathogen within a facility in the ability of that pathogen to persist over time. The parameter classes in the matrices, in order, are: doctor, nurse, patient, equipment, surfaces.

**Fig. 3.**
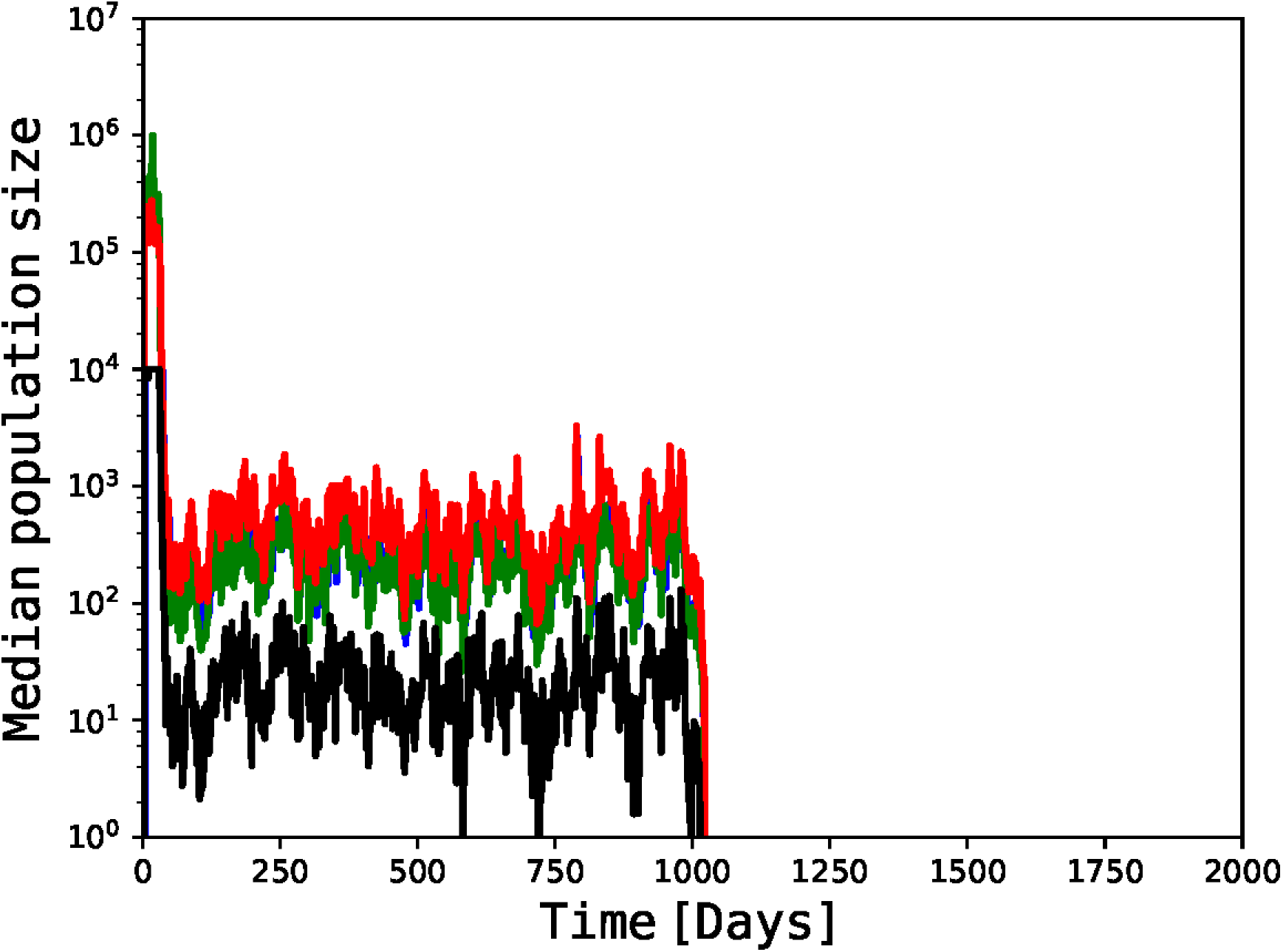
Median size of patch populations over time for 1,000 computational simulation trials. After the initial forcing period, median patch population size fluctuates but remains at a low level for extended periods of time, here over 1,000 days. The metapopulation persists at an equilibrium between patch population births and deaths, with the rates of each affected by transmission and cleaning. At any point during this time period, a jackpot even could occur which results in a secondary active infection that would start a new forcing period. Median population size for surface patches (in yellow, not visible) remains close to 0 throughout timeframe. After approximately 1,000 days, the other patch type populations also persist at extremely small median sizes. Median time to metapopulation extinction in our simulation trials was 506 days.

**Fig. 4.**
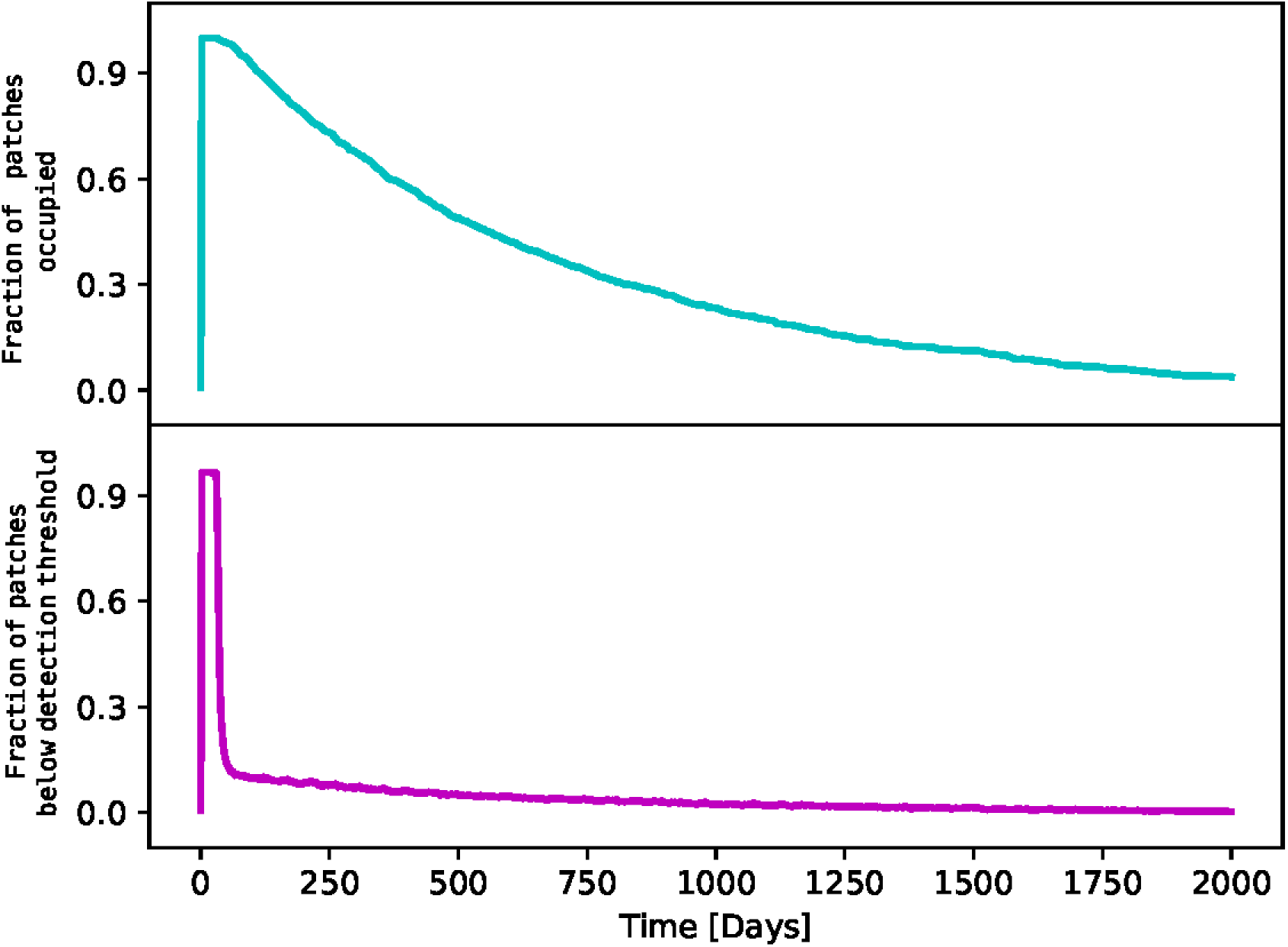
Fraction of occupied patches over time as compared to fraction of populations above threshold of detectability. A comparison of the average fraction of patches occupied, regardless of local population size, and the fraction of patches with local population sizes greater than the threshold of detectability, here set to 10^3^ cells/patch. The slow decline in metapopulation (first panel) suggests that hospital pathogens survive for much longer periods of time than previously suspected without a high probability of detection through most monitoring protocols. For instance, at 500 days, approximately 50% of all patches have at least some pathogens in them, but only about 15% have more than the number required to be detected with any reasonable probability.

At the beginning of our simulations, all islands are free of pathogens. We introduce a pathogen on a patient for a given amount of time, 30 days (t_start_), after which this index patient is removed. We run our simulations by randomly drawing, every day, a sequence of interactions between islands and of cleaning events, and by following the changes in the growth and locations of the populations of pathogens over time.

## Results

A total of 1,000 Monte Carlo trials were conducted. The overall number of trials was determined by verifying that the number be sufficient to ensure that our average results do not depend on the total simulations run. We simulate a situation where an index patient carrying a pathogen is admitted to a naïve hospital ward for t_start_=30 days. Sensitivity analyses indicate that all results are robust to two-fold increases/decreases to interaction matrix (*N_ij_*) values (see SI for more information).

All simulations begin with a single index patient. Trials show all patches quickly reaching maximum carrying capacity and remaining at those levels for the duration of the infected patient’s stay. Following removal of the source population, island population sizes decline, but do not go extinct for extended periods of time. This results in the continued transmission of the pathogen between hospital workers, environmental patches, and novel patients during that period. Without a new source population (second index patients were not introduced in trials), simulations indicate that the metapopulation will eventually go extinct, but the time to total metapopulation extinction in the model is a function of the frequency and level of cleaning events affecting individual patch populations.

This is the first of several unexpected results of our model simulations. The longer a pathogen metapopulation can remain extant within a hospital, the greater the probability of a secondary infection occurring in a susceptible patient. In our model, any secondary infection represents a stochastic jackpot event in which a limited number of cells founds what will become a large new island population with which other potential hosts (short and long term) are expected to interact. When such a secondary infection occurs, patch populations quickly reach carrying capacity once again, thereby resetting the time to extinction clock for the metapopulation as a whole. As a result, the metapopulation can remain in existence indefinitely. From an epidemiological perspective, there need not be any direct contact at all between the primary and secondary infected patients, so long as the time between the two cases is less than the time to metapopulation extinction and there exists a transmission path, regardless of how indirect, between the two infected patients. The reason for the extended persistence of the metapopulation is that in order for permanent extinction of the metapopulation to occur, all islands must be extinct at the same moment. If any island population remains, regardless of the number of individuals, the possibility remains for other islands to be repopulated over time and for additional infections to occur. In this manner, the model explains the discontinuity of infections often observed in hospital outbreaks.

A second outcome of model simulations relates to the small size of island populations during the period between loss of the source population and extinction. Model parameterization set limits to patch population sizes based upon island class. These limits are in line with empirical data currently available ^6,26,37,^38^^. In most simulation trials, the size of patch populations quickly dropped below carrying capacity once the source population was removed. The island populations remained extant at small sizes for significant periods of time before going extinct. However, the local populations continue to survive during this time, even at such small scales. Patches surviving at levels indicated by our simulations suggest that many of these islands would be too small to be readily detected by most pathogen sampling techniques currently in use in healthcare facilities.

Typically, hospitals employ culture-based protocols for the presence/absence detection of pathogens. However, all such methods have a lower limit to their resolution of detection ^2,39^. In large part, this limit may be due to sampling itself: in order to detect a pathogen, staff must first swab then culture a strain. Successful detection is impacted by, among other things, the likelihood of sampling enough of the relevant bacteria (typically from a patient). If the same techniques are applied for surveillance of staff and environment during which island populations exist in small populations, the probability of detection is greatly diminished, leading to a large false negative bias. In fact, our simulations suggest that metapopulations can persist on islands for extended periods below a reasonable size threshold of detectability, providing faulty data upon which facility personnel may base critical decisions.

An additional result indicates that a metapopulation model, while robust to changes in interaction or transmission parameters, is sensitive to the number of local populations that make up the metapopulation system. For patch classes such as doctors, nurses, and the patients themselves, the number of possible patches is relatively limited: most pathogen islands will be on hands and most of these will have high frequencies of cleaning with high effectiveness. However, the number and duration of non-human islands, such as those on equipment and surfaces are more problematic. The model here is sensitive to the number of these islands, with minimum thresholds being vital for metapopulation survival (Fig 5). This may present a unique target for HAI control measures if reliable methods for detecting and monitoring such patches can be implemented in clinical environments. The present simulations all utilized a conservative number of 4 surface patches per patient. A distinctly larger number of surfaces can be envisioned, including sites known to be easily contaminated such as bedrails, door knobs, toilet handles, call buttons, faucet, etc., suggesting that the median time to extinction for pathogen metapopulations in real-world environments is even longer than that suggested in our simulation trials.

**Fig. 5.**
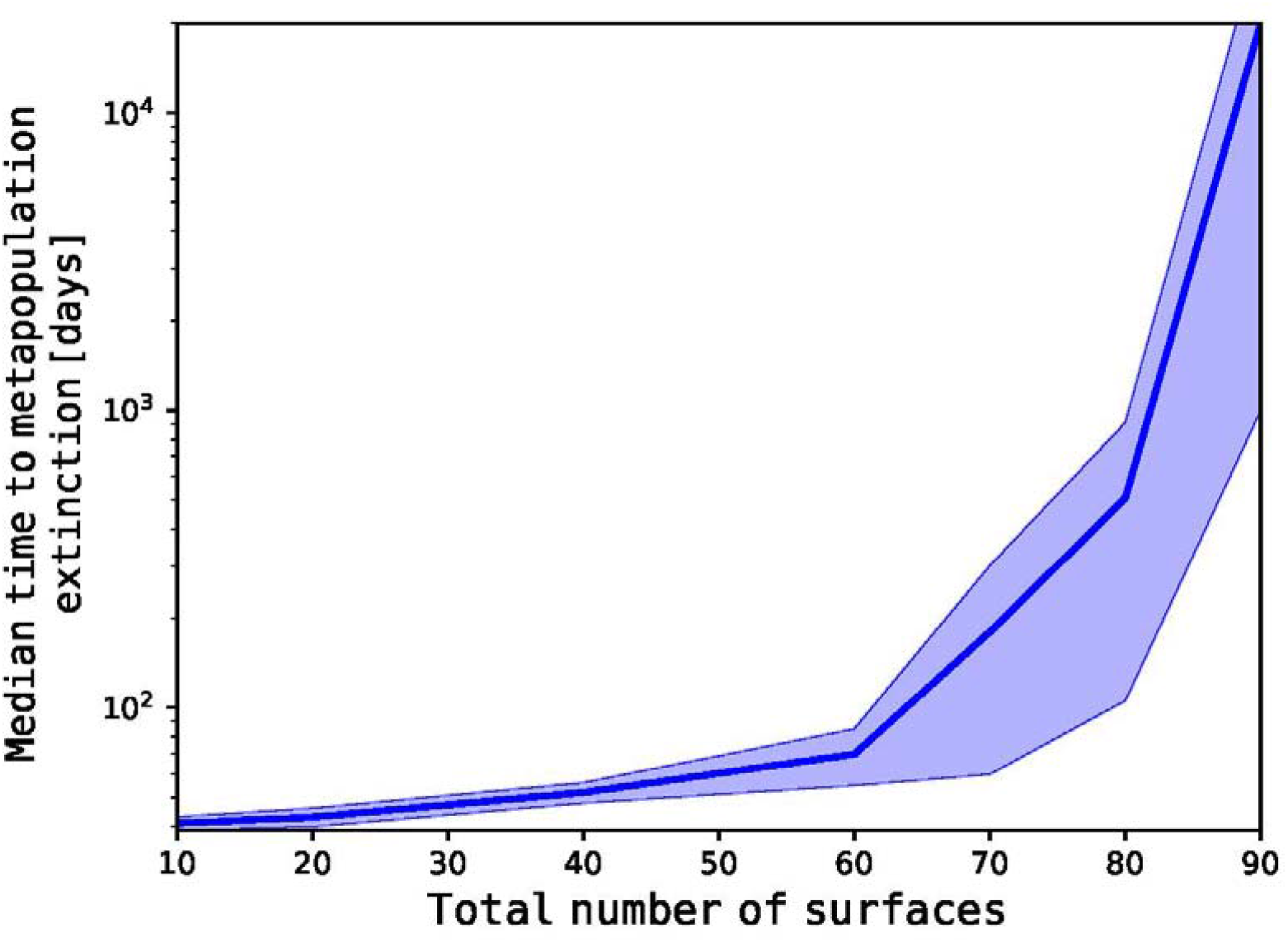
Importance of the Number of Surface-type Patches. While sensitivity analyses indicate the model to be robust to interaction or transmission values, the total number of patches within a metapopulation is important to the system. Here, the amount of available patches on surfaces must be greater than 60 in order for the metapopulation to resist relatively rapid extinction. The simulations here concern a total of 20 patients and 80 surface patches resulting in only 4 patches per patient. A larger number of patches per patient can easily be envisioned. The possibility of threshold values existing provide possible intervention targets that can be exploited for a hospital ward based upon the parameters specific to that clinical site. However, such an approach would require accurate and regularly updated information about the ward. Note the Log scale on the Y-axis. Shaded area indicates ±1 SD.

In addition to the sensitivity of the model to patch number, also noteworthy is the model’s robustness to the interaction and transmission levels that define the system. By having low sensitivity to changes in these parameters (denoted in *N_ij_* and *P_ij_*), simulation trials suggest that a stable equilibrium can be reached for a wide range of physical constraints across hospital conditions. This results in a resilient population-level system by which pathogens can survive and adapt to many different specific healthcare facilities so long as the core structure and function of the hospitals are alike across sites. The outcome is that hospitals are susceptible to endemic metapopulations of pathogens by their very design and function.

## Discussion

Mathematical models are often used as a mechanism by which we can understand the dynamics of infection transmission and spread in hospital environments ^33^. While models have proven helpful in understanding how HAIs spread, they can be only as accurate as the biology that underlies them. Here we propose a model in which pathogen populations in hospital environments act as metapopulations, with small patch populations interconnected by migration between ephemeral islands. The size of these island populations, their duration, and the migration between them are dictated by human behaviors, many of which are known to affect HAI transmission. While each aspect of our model has been observed in isolation ^2,5,6,40^, to our knowledge, they have never been integrated into a common framework such as presented here. The integration of these data into a cohesive model will allow for a more complete understanding and analysis of pathogen transmission in hospitals.

Our simulations indicate that by existing in a metapopulation structure, pathogens can survive within a hospital for extended periods of time even without a source population. In essence, each patch population contributes to a diffuse reservoir that resists extinction through a bet-hedging strategy in which the metapopulation is never in any one place at any single time. This results in the metapopulation of pathogens remaining capable of causing secondary infections long after the source is removed, explaining one of the troubling aspects of HAIs. Further, local populations evade detection by hospital pathogen surveillance protocols in part by existing in unexpectedly small groups over diverse locales. The scale of applicability for this model is yet to be determined and may be specific to the unique dynamics of particular healthcare environments. Next steps require informing the model with parameter values specific to individual hospitals to understand the physical limits of metapopulations in specific institutions (individual rooms, wards, entire wings, etc.). Our model invites study of several aspects of hospital pathogen population biology, with the potential to guide development of novel intervention protocols and may also provide beneficial insight into the ecology and evolution of infectious disease and how disease causing agents persist in human dominated environments.

## Acknowledgements

FS and PC thank Columbia Frontiers of Science for support as well as fostering opportunities for interdisciplinary collaboration. This work was supported by Columbia Frontiers of Science Fellowships to FS and PC. The authors declare no competing interests.

